# Bayesian priors for tree calibration: Evaluating two new approaches based on fossil intervals

**DOI:** 10.1101/014340

**Authors:** Ryan W. Norris, Cory L. Strope, David M. McCandlish, Arlin Stoltzfus

**Affiliations:** Department of Evolution, Ecology and Organismal Biology, Ohio State University, 4240 Campus Dr., Lima, OH 45804,USA; Bioinformatics Research Center, Box 7566, North Carolina State University, Raleigh, NC, 27695, USA; Department of Biology, University of Pennsylvania, Philadelphia, PA 19104, USA; National Institute of Standards and Technology, Institute for Bioscience and Biotechnology Research, 9600 Gudelsky Dr., Rockville, MD, 20850, USA

**Keywords:** Ghost Lineage Length, Molecular clock, Penultimate Gap, Time-tree, phylogeny calibration

## Abstract

**Background:** Studies of diversification and trait evolution increasingly rely on combining molecular sequences and fossil dates to infer time-calibrated phylogenetic trees. Available calibration software provides many options for the shape of the prior probability distribution of ages at a node to be calibrated, but the question of how to assign a Bayesian prior from limited fossil data remains open.

**Results:** We introduce two new methods for generating priors based upon (1) the interval between the two oldest fossils in a clade, i.e., the penultimate gap (PenG), and (2) the ghost lineage length (GLin), defined as the difference between the oldest fossils for each of two sister lineages. We show that PenG and GLin/2 are point estimates of the interval between the oldest fossil and the true age for the node. Furthermore, given either of these quantities, we derive a principled prior distribution for the true age. This prior is log-logistic, and can be implemented approximately in existing software. Using simulated data, we test these new methods against some other approaches.

**Conclusions:** When implemented as approaches for assigning Bayesian priors, the PenG and GLin methods increase the accuracy of inferred divergence times, showing considerably more precision than the other methods tested, without significantly greater bias. When implemented as approaches to post-hoc scaling of a tree by linear regression, the PenG and GLin methods exhibit less bias than other methods tested. The new methods are simple to use and can be applied to a variety of studies that call for calibrated trees.

## Background

Time-scaled phylogenies, that is, phylogenies calibrated so that the branch lengths are proportional to time, are vital tools in studies of macroevolution (Slater and Harmon, 2013). Typically, time-calibrated phylogenies are generated by a phylogeny inference method applied to molecular sequence data, with fossil dates assigned to designated nodes as calibration points. Recent "tip-dating" methods (Pyron 2011; Ronquist, Kloppstein, et al. 2012) represent a distinctive new approach. By including a matrix of character data on fossils, such methods allow fossil taxa to be treated as terminals (tips) and included directly with extant taxa (and their sequence data) in a unified approach to inference that avoids the need to make prior assignments of fossils to nodes. Unfortunately, tip-dating approaches require coded morphological matrices for both extant and extinct taxa, unavailable in many, if not most, situations (Heath et al. 2014)

In the vast majority of studies, each fossil is assigned to a clade, and the corresponding fossil dates are integrated as constraints or priors on the age of the clade. Yet, a persistent problem has been to understand what precise constraint or prior is implied by the assignment of a fossil to a clade. Analyses that employ a molecular clock often use fossils as point estimates of divergence times (Sarich and Wilson 1967; Baldwin and Sanderson 1998; Near, Bolnick, et al. 2005; Near, Meylan, et al. 2005). In reality, the fossil dates assigned to a clade represent a minimum age estimate (Marshall 1990; Conroy and Van Tuinen 2003; Magallón 2004; Forest 2009). A variety of methods are based on this logic of the "hard minimum" (Sanderson 1997; Thorne et al. 1998; Yang and Rannala 2006; Benton and Donoghue 2007; Marshall 2008). In particular, in the approach of Marshall (2008), an uncalibrated ultrametric tree is calibrated separately with each fossil date, and the fossil that yields the oldest root estimate is selected as the single best calibration point.

Currently there are multiple software implementations of Bayesian divergence time estimation with a relaxed molecular clock (MCMCtree by Yang 2007; BEAST by Drummond et al. 2012; and MrBayes by Ronquist, Teslenko, et al. 2012). These sophisticated tools allow the user to impose, not merely a hard constraint on the age of a node, but a prior probability distribution. However, an open question is how to use fossil data to derive a prior that is objectively justifiable (Ho 2007; Ho and Phillips 2009). While a correctly assigned and dated fossil may serve as a hard minimum on a node's age, little is known about how to infer the shape of the rest of the distribution, even though the shape of the prior can have a substantial impact on results (Inoue et al 2010; Warnock et al. 2012). That is, the availability of generalized Bayesian software has not solved the problem of fossil-based phylogeny calibration, in the absence of valid and practical methods to assign informed priors.

Here we present two new methods to assign priors that have a known shape based on an explicit model, that allow multiple calibration nodes, and that can be implemented easily in BEAST (Drummond and Rambaut, 2007). The use of multiple calibrations has been shown to minimize error (Duchêne et al. 2014). Both of our new approaches follow from the logic of Solow (2003; see also Robson and Whitlock, 1964; van der Watt 1980) regarding intervals between known fossils assigned to a clade. That is, they are not based on single fossils, but on pairs of fossils that specify an interval. One useful interval is the gap between the two oldest known fossils assigned to a clade, here called the “penultimate gap” or **PenG**. The other interval of interest is the gap between the oldest fossil on one lineage, and the oldest fossil on a sister lineage, the “ghost lineage length” or **GLin**. Under the conditions we define, PenG and GLin/2 are estimates of the ultimate gap between the oldest known fossil and the true origin of the clade. Furthermore, we show (using the Jeffreys prior for the unknown rate of fossil deposition and discovery) that given PenG (or GLin) the length of the ultimate gap follows a log-logistic distribution with a shape parameter of 1 and a scale given by PenG (or GLin/2). This distribution can then be closely approximated by a lognormal distribution where the underlying normal distribution has a mean of log PenG (or log (GLin/2)) and a standard deviation of 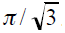.

Approaches to calibration based on PenG and GLin can be implemented easily using the lognormal approximation in BEAST, a widely used software package (Drummond and Rambaut 2007). We evaluated the performance of these 2 approaches relative to several other approaches using fossil dates (Marshall, 2008; Dornburg et al., 2011). More specifically, we simulated molecular and fossil data using 3 different tree shapes of an 8-taxon tree, under low, medium and high rates of fossil recovery. We then conducted a comparative evaluation of the accuracy of methods for (1) post-hoc calibration of the true tree, (2) post-hoc calibration of a tree with branch lengths inferred from simulated sequences, and (3) Bayesian inference of a calibrated tree combining sequences with priors based on fossil data. When implemented as approaches to post-hoc calibration, the PenG and GLin methods exhibit less bias than the method of Marshall (2008), or the method of using fossil ages as point estimates of node ages. When implemented as approaches to assigning priors for the inference of a scaled tree in BEAST, the PenG and GLin methods are more precise than the other methods tested, without significantly greater bias, resulting in inferred divergence times that are closer to the true times. Compared to other approaches for using limited fossil data, the new interval-based methods appear to be nearly as simple and widely applicable, while being more accurate and rigorous.

## Methods

### New approaches

Figure 1 shows a hypothetical node representing the most recent common ancestor of two sister taxa, A and B, with fossil information relevant to estimating their divergence time. We assume that fossils are dated precisely and display characters that allow them to be correctly assigned. Although fossil taxa are more appropriately conceived as belonging to side branches (Foote 1996), we have collapsed these side branches, assigning fossils directly to lineages. Implications of these and other simplifying assumptions are addressed in the Discussion.

**Figure 1.**
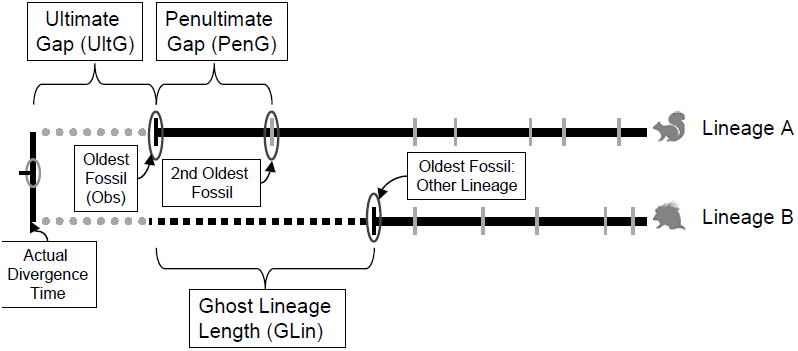
An illustration of terms used in the text. Shown is a part of a phylogeny to be calibrated, including a node (left) with 2 daughter lineages (right), A and B. Hatches indicate dated fossils (collapsed onto main branches for simplification). The observed fossil range is shown in solid black. A ghost lineage of known length, GLin, is present in lineage B, as the two lineages diverged at the same time, but lineage A has an older fossil. The actual divergence time is older than Obs, the age of the oldest fossil. The difference between Obs and the actual age is UltG, the ultimate gap. The difference between the two oldest fossils, PenG or the penultimate gap, is a point estimate of UltG. Note that the two oldest fossils assigned to a clade might appear on the same or different lineages, but only the former possibility is illustrated in this figure.

For our purposes, the observed (abbreviated **Obs** hereafter) divergence date is defined as the age of the oldest fossil attributable to either lineage that descends from their common ancestral node. This value will almost always underestimate the actual divergence time. We refer to the difference between the true divergence date and Obs as the ultimate gap, abbreviated **UltG** hereafter.

We focus here on two potential estimators of UltG, although either gap analysis (Strauss and Sadler 1989; Marshall 1994, 1997) or collector’s curves (Kalmar and Currie 2010) hold potential as alternate approaches. The first estimator is the interval between the age of the oldest fossil and the age of the second oldest fossil, regardless of lineage. We call this difference the penultimate gap or **PenG** (Fig. 1). Solow (2003) and Robson and Whitlock (1964) argue for using PenG as an estimate of UltG. The second estimator is the interval between the ages of the oldest fossil in each of two descendent lineages. Müller and Reisz (2005) listed agreement in age for first-appearance dates between daughter lineages as one of their three criteria of a good calibration: the richer the fossil record, the better the agreement. Following Müller and Reisz (2005), we refer to this interval as the ghost lineage length or **GLin** (Fig. 1).

In order to understand the relationship between PenG, GLin and UltG, we introduce some additional notation, as well as mathematical assumptions. First, let us name two descendent lineages A and B, as in Figure 1. We suppose that fossils destined to be observed appear along each of lineages A and B according to two independent, inhomogeneous Poisson processes. In particular, let us write *λ*_*A*_(*t*) as the rate at which fossils appear on lineage A at time *t* and *λ*_*B*_(*t*) as the corresponding rate for lineage B, where we have set the actual divergence time to be *t* = 0. In order for our method to work, we must assume that fossils occur frequently enough and that these rates change slowly enough that they are approximately constant from the actual divergence time through the time period where the first several observed fossils are likely to occur. We thus ignore the time-dependence of the rate s and treat them as unknown constants, which we write as *λ*_*A*_ and *λ*_*B*_.

To see why PenG and GLin are informative with regard to UltG, it is helpful to consider the expectations of these times. For the case of PenG, we need only consider the times at which fossils appear, and not which lineage each fossil appears on. Because fossils occur as independent Poisson processes with rate *λ*_*A*_ on lineage A and *λ*_*B*_ on lineage B, the appearance of fossils without regard to lineage is a Poisson process with rate *λ* = *λ*_*A*_ + *λ*_*B*_. Furthermore, because fossils appear as a Poisson process with rate *λ*, if we begin watching the process at any given time *t*, the waiting time until the next fossil arrives is exponentially distributed with rate *λ*. If we begin watching at *t* = 0, this waiting time is simply UltG, and if we begin watching at Obs, the waiting time is PenG. Thus UltG and PenG are both exponentially distributed with rate *λ* and their expected values are both equal to 1/*λ*.

In order to relate GLin to UltG, we assume further that; *λ*_*A*_ = *λ*_*B*_ over the scale of the interval in question (the initial divergence of A and B represented by the oldest fossils in the clade). Under this assumption, the first fossil is equally likely to appear on either lineage. The waiting time for the next fossil to appear on the other lineage is then exponential with mean 2/*λ*, since fossils appear on any one lineage as a Poisson process with rate *λ* / 2. Because UltG is exponential with rate *λ*, one can see that the expected duration of GLin is twice as long as UltG.

The above arguments suggest using PenG and GLin/2 as point estimates of UltG. However, we are also interested in using the observed values of UltG and GLin to derive an objectively justifiable prior distribution for UltG for use in phylogenetic inference. To this end, we will take a Bayesian approach.

Our first task is to choose a prior for *λ*, the total rate that fossils appear across both lineages. Because we would like our method to be based on the fossil data alone and not incorporate other information, we will take the prior on *λ* to be the uninformative, improper prior *p*(*λ*) = 1/*λ*. This is the Jeffreys’ prior for the rate of an exponential random variable and corresponds to a uniform distribution on all possible values of log *λ*. Now that we have chosen a priorfor *λ*, we need to update this prior based on our observed value of PenG. In particular, suppose we observe PenG = *x*. Because a gamma distribution is the conjugate prior of the likelihood function for the rate of an exponential random variable and *p*(*λ*) = 1/*λ* corresponds formally to the probability density function of a gamma distribution with its shape and rate parameters set to zero, the posterior distribution for *λ* given *x*, *p*(*λ* | *x*), is gamma-distributed with shape 1 and rate *x*, i.e., it is exponential with rate *x*.

Given this posterior distribution on the total rate at which fossils occur in either lineage, we then wish to derive a posterior distribution for UltG. Supposing that the true value for UltG is y, we can write the probability density for the posterior distribution for UltG given PenG as *f*(*y* | *x*). This density can then be found by treating *λ* as a nuisance parameter, and, in particular, we have:

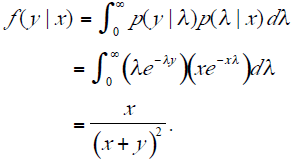

The posterior distribution for UltG is thus a special case of the log-logistic distribution where the shape is 1 and the scale is *x* (it is also a special case of the generalized Pareto distribution with location 0, shape 1, and scale *x*). It is worth noting that while this distribution is quite heavy tailed (its expectation is undefined), the median of a loglogistic distribution is equal to its scale, which in this case is our observed value for PenG. Thus, the median of our posterior distribution for UltG is equal to the point estimate we derived earlier.

A similar argument can be used if we observe GLin = *z*, in which case we must again assume *λ*_*A*_ = *λ*_*B*_. Given GLin = *z* and, assuming that the prior for *λ* has density prior *p*(*λ*) = 1/*λ*, the posterior distribution for *λ* is exponential with rate *z*/2. Then, treating *λ* as a nuisance parameter, the posterior probability density function for UltG is given by

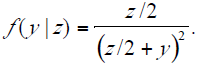

This distribution is also log-logistic, with shape 1 and scale *z*/2.

To summarize, given an observed value for PenG or GLin and an uninformative prior *p*(*λ*) = 1/*λ* on the rate of fossil occurrence, we can compute a posterior distribution for UltG. This posterior distribution for UltG can then be used to construct a prior for the actual divergence time for use in subsequent phylogenetic analysis.

It is important to understand that the priors produced by this method, while strongly informative regarding the minimum age of a node, are only weakly informative with regard to the maximum. To see what is going on, note that our prior for the rate of fossil occurrence *p*(*λ*) = 1/*λ* corresponds to the improper, uninformative prior *f*(*y*) = 1/*y* for the value of UltG. We are updating this uninformative prior with a single observation and thus the resulting posterior distribution for UltG is only weakly informative (e.g. it has infinite expectation, and so its higher central moments are undefined). However, this uninformativeness is entirely appropriate: given limited fossil data, we only have very limited information about the rate of fossil occurrence, and therefore limited information about the true value of UltG. The goal here is to accurately and objectively summarize our degree of ignorance.

### Simulated datasets

Having developed a framework for constructing priors based on PenG and GLin, we can compare them with other approaches, using simulated data. We compared our new methods to one another, to trees calibrated using Obs without correction, and to trees calibrated using Marshall’s (2008) method. Marshall’s (2008) approach, abbreviated **M08** hereafter, begins with an uncalibrated ultrametric tree. Each potential fossil calibration is used individually to estimate the age of the root of the tree. The calibration that yields the oldest root estimate is selected as the single optimal choice for calibrating the tree.

Using a script in R (R Development Core Team 2011), random node ages were generated for 8-taxon trees with three increasingly asymmetric branching patterns: fully symmetric, intermediate, and fully pectinate (ladder-like), where the intermediate tree has the topology (((((A,B),C),D),(E,F)),(G,H)). This approach affords a set of trees for which downstream analysis can be efficiently performed, as clade composition is consistent within each tree shape, though it may not result in the most biologically realistic tree shapes (Raup and Gould 1974; Guyer and Slowinski 1991; Stadler 2010).

All trees were ultrametric and had a root age of 100 Ma. Node ages were obtained by generating a set of *n* random numbers less than 100 where *n* is the largest number of nodes available along any path to the root (2, 4, and 6 values for symmetric, intermediate, and pectinate respectively) and assigned in order by age. Remaining nodes were assigned ages by generating a random number less than the previously assigned ancestral node age.

Trees were provided as input to a modified version of indel-Seq-Gen version 2.2.03 (Strope et al. 2009), which assigned random fossil dates along branches and generated 1000 bp of DNA sequence under a generalized time reversible (GTR) model of evolution and a strict molecular clock. The parameters of the GTR model were based on actual data from a set of pooled nuclear and mitochondrial genes in mammals (Norris 2009) to ensure that realistic values were applied. Fossil sampling occurred as a Poisson process along each branch on the tree and was repeated at three rates: one fossil per 4 Ma, 10 Ma, or 25 Ma. For a given tree, molecular data were only used from 4 Ma runs and discarded for 10 Ma and 25 Ma runs in order to save computer time and to facilitate the direct comparison between a given set of molecular data but under different fossilization rates. A total of 1000 runs were conducted for each combination of tree and fossil sampling rate. In all, 9000 runs of indel-Seq-Gen were conducted on 3000 input trees.

We compared the effects of different approaches to calibration in three ways: (1) post-hoc calibration of the true (input) tree, (2) post-hoc calibration of the maximum clade credibility (MCC) ultrametric tree inferred from the simulated molecular data in BEAST in the absence of prior calibration, and (3) inference of the divergence dates in BEAST using the molecular data with fossil calibrations included as prior distributions. This three-tiered approach was designed to tease apart (1) biases inherent to the way different approaches use fossil dates, in the absence of error introduced by tree inference, (2) biases emerging in the context of error in tree inferences, and (3) biases and precision when calibration priors interact during a full Bayesian analysis.

For the post-hoc calibrated true tree test, the true (input) tree was converted to an uncalibrated ultrametric tree by dividing all nodes by the root height (100 Ma). This ultrametric tree was then calibrated using Obs, M08, PenG, and GLin information. For the Obs method, the Obs value was assigned as the calibration at each of the 7 nodes on the tree. These fossil dates were compared to the relative ages on the ultrametric tree using linear regression with the intercept constrained at 0 (Conroy and van Tuinen 2003). The resulting equation was used to calculate the age at all nodes. This approach allowed for the tree to be calibrated based on the central tendency of all potential fossil calibrations.

The M08 method assigns the calibrating node as the node that yields the oldest age estimates across the tree, consistent with equation 11 in Marshall (2008). We applied the absolute age of this fossil as a calibration, and calculated the ages of all nodes. Our M08 estimates therefore correspond to Marshall’s (2008) minimum age bracket; we did not calculate an upper confidence interval.

The PenG and GLin methods that we propose here are intended as methods for assigning multiple Bayesian priors simultaneously: in order to carry out the quite different task of post-hoc calibration (in the first two tests), we implemented simplified PenG and GLin approaches in which the input tree is scaled by linear regression using the Obs + PenG or Obs + GLin/2 values, respectively. The resulting methods are strictly comparable to an existing method that we refer to as Obs, in which the input tree is scaled by linear regression using all available Obs values as point estimates of node ages (see Conroy and van Tuinen 2003).

For the PenG method, the value for PenG was calculated as the difference between the two oldest fossils from daughter lineages. The values for Obs + PenG were compared to the relative ages on the ultrametric tree using linear regression as described above and the resulting equation was used to calculate the age at all nodes. Note that for the post-hoc calibrated true-tree test, PenG is treated as a point estimate instead of a distribution.

For the GLin method, the value for GLin was calculated as the difference between Obs and the oldest fossil on the other daughter lineage at a node. The values for Obs + GLin/2 were compared to the relative ages on the ultrametric tree using linear regression as described above and the resulting equation was used to calculate the age at all nodes. Again, GLin/2 is treated as a point estimate for this test.

In our analysis, calibrations for a particular method are limited (appropriately) by stochastic variation in the simulated fossil record. The number of simulated fossils assigned to a branch is a random variable that may take the value of zero: when the lack of fossils renders undefined the value of GLin, PenG, or Obs for a node, the calibration simply proceeds on the basis of any other nodes for which suitable data are available. Bias was calculated at each node on the tree by subtracting the actual age (from the initial input tree) from the estimated age. Positive bias indicates that the estimates for node age are older than their actual age while negative bias indicates an estimate that is too recent. Mean bias was calculated for each tree, and this value was stored. This process was automated using a script in R, and repeated across all 9000 simulated fossil datasets. We tested each method to determine if bias values were significantly different from 0, which indicates no bias, using a Wilcoxon signed rank test in R. Bias was compared across methods using a Friedman’s test with post-hoc analysis in R.

For the post-hoc calibrated inferred tree test, the molecular data were used to generate an uncalibrated ultrametric tree in BEAST (version 1.6.1, Drummond and Rambaut 2007). The age of the root of the tree was forced to 1.00 with a normal prior with mean = 1.0 and standard deviation = 0.0001, which yields results that round to 1.0 within 3 decimal places. Data were analyzed using a GTR model, an uncorrelated lognormal relaxed clock, and a Yule speciation prior. Output tree files from BEAST were summarized in TreeAnnotator (version 1.6.1) and the maximum clade credibility (MCC) tree was saved.

The MCC tree was then calibrated using the approaches outlined above for the actual tree using Obs, M08, PenG, and GLin methods. Again, bias was calculated for each node on the tree, and the mean of these bias values was stored. This process was repeated for the first 100 trees of each shape for each fossilization rate for a total of 900 fossil datasets. We tested each method to determine if bias values were significantly different from 0 and the bias present in different methods was compared as described above.

For the final test, we evaluated the approaches to calibration when priors are explicitly set as part of a Bayesian analysis. Normal priors with a standard deviation = 0.0001 were used for Obs and M08 methods, making them essentially fixed points. BEAST has no option for a log-logistic or generalized Pareto distribution, but a lognormal distribution where the underlying normal distribution has a standard deviation 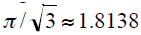, is a very close approximation— derived by setting the mean and variance of the normal distribution (underlying the lognormal distribution) equal to the mean and variance of the logistic distribution (underlying the log-logistic distribution). This approximation is good for values up to about 100 times longer or 100 times shorter than PenG (or GLin/2) but underestimates the probability of even more extreme deviations from our point estimate (Fig. S1). To use this approximation in BEAST, the corresponding line in the XML input file should be set to <logNormalPrior mean="log(x)" stdev="1.814" offset="Obs" meanInRealSpace="false"> where ‘Obs’ is the value of Obs, and x is the value of PenG (PenG method) or GLin/2 (GLin method).

To adapt M08 for a Bayesian analysis, we choose the single calibrating node via Marshall's method, and run the BEAST analysis using that single fossil as a calibration. We designate this approach **M08^*^**. In our application of the M08^*^ method, only the calibrating node is fixed and all other nodes are estimated. Among these estimates are values that are younger than the oldest fossil at non-calibrating nodes, a situation Marshall (2008) explicitly sought to avoid. M08 was designed to be used *a posteriori* on an uncalibrated tree inferred from genetic data, while our challenge was to adopt an approach that would test the method when the calibration was applied *a priori*. A common approach in the literature is to set minimum age constraints whenever any fossil information is present along a daughter lineage. We implemented this approach (which we abbreviate **M08min**) in BEAST by adding a uniform prior with a minimum value = Obs and a maximum value = 1000 for all nodes except the one already assigned as a calibration in M08^*^.

We also added an abbreviated version of the method of Dornburg et al. (2011), which we call **aD11**. Concerned with Marshall’s (2008) reliance on a single calibration and a single ultrametric tree, Dornburg et al. (2011) developed an approach that selected the subset of potential fossil calibrations that were not significantly different from the single calibration that yielded the oldest age estimate. All of the selected fossil calibrations are then used in a final Bayesian analysis with prior shapes chosen so that the upper 95 % confidence interval matches equation 11 from Marshall (2008). Although we were interested in evaluating the performance of this method, its full implementation requires an infeasible number of BEAST runs, as many as there are potential fossil calibrations (up to 7 per run in our case).

To reduce the computational burden, we took advantage of the fact that Marshall’s (2008) equation 11 is identical at all variables for all calibrating nodes except for being scaled by the age of the calibration itself. Based on this situation, we adopted an approximation of the Dornburg et al. (2011) method that would require only a single initial BEAST analysis by calibrating the root of the tree with a lognormal prior where zero offset = 1.0, standard deviation = 0.5 (as used by Dornburg et al. 2011), and the mean was set so that the upper 95 % of the distribution matched Marshall’s (2008) equation 11 when the calibrating fossil is fixed at 1.0. The primary calibrating node was selected by scaling (as per equation 6 in Marshall 2008) the MCC tree generated by TreeAnnotator, and upper and lower bounds on this scaling factor were calculated by scaling the ends of the 95 % HPD estimate of values at that node. The 95 % HPD range also was scaled for all other nodes with fossils, and the other calibrating nodes for the final analysis were identified as those whose empirical scaling factor values overlapped the values for the primary node. The final BEAST analysis included the selected calibrations with lognormal priors where zero offset = Obs, standard deviation = 0.5, and with the mean set so that the upper 95 % of the distribution matched Marshall’s (2008) equation 11.

We conducted a brief test comparing our abbreviated version (aD11) to the full version of Dornburg et al.’s (2011) method. We applied the complete Dornburg et al. (2011) method to 3 simulation runs using 3 different fossilization rates on 3 different simulated trees of the intermediate topology (see above). In all instances, our aD11 method selected the same set of node and fossil combinations as the complete Dornburg et al. (2011) approach. Specifically, both methods chose to calibrate using the fossils at all 7 nodes in the high fossilization (every 4 Ma) example, using fossils at 5 nodes in the medium fossilization (every 10 Ma) example, and using fossils at 6 nodes in the low (every 25 Ma) example. We believe aD11 will exhibit similar patterns to Dornburg et al. (2011), but do not claim that aD11 will lead to the same results in all cases, thus we do not claim to have tested their method fully.

BEAST runs were repeated as outlined above and bias was calculated as stated earlier. In addition to bias, we also were interested in the accuracy and precision of each method. All nodes were evaluated to see if the actual age was included in the 95 % HPD; if so, the width of the 95 % HPD was recorded. We tested accuracy by determining whether the actual age at a node fell within the 95 % HPD estimate and the length of that interval is our evaluation of precision. Friedman’s test with post-hoc analysis was used to evaluate whether accuracy differed among methods. Tukey’s HSD (Honestly Significant Difference) test was performed to determine whether methods yielded significantly different 95 % HPD lengths for accurate nodes. The percentage of accurate nodes was also calculated; a well-calibrated Bayesian analysis should be accurate 95 % of the time, matching the 95 % HPD (Dawid 1982). This process was repeated for the first 5 trees of each shape for each fossilization rate for a total of 45 fossil datasets. To test the 5 methods, a total of 225 BEAST analyses were conducted at this stage in addition to the extra analyses required to set values in aD11 analyses. Input trees, simulated fossil and genetic data, BEAST .xml input files and TreeAnnotator output trees are available (http://github.com/arlin/PenG_and_GLin).

## Results

The results of comparing methods for post-hoc calibration of the true tree (Figure 2 and Table 1) indicate that PenG and GLin have the least bias, though all four methods exhibit significant (albeit sometimes slight) underestimation bias. Figure 2 shows box plots displaying the distribution of the mean bias (estimated age minus actual age) of nodes on individual trees. The bias for the Obs method is significantly larger than the M08 method, which is in turn significantly larger than PenG or GLin methods. The PenG method significantly outperforms all other approaches at high (every 4 Ma) and low (every 25 Ma) fossilization (Table 1), but is not significantly better than the GLin method at medium fossilization (every 10 Ma).

**Figure 2.**
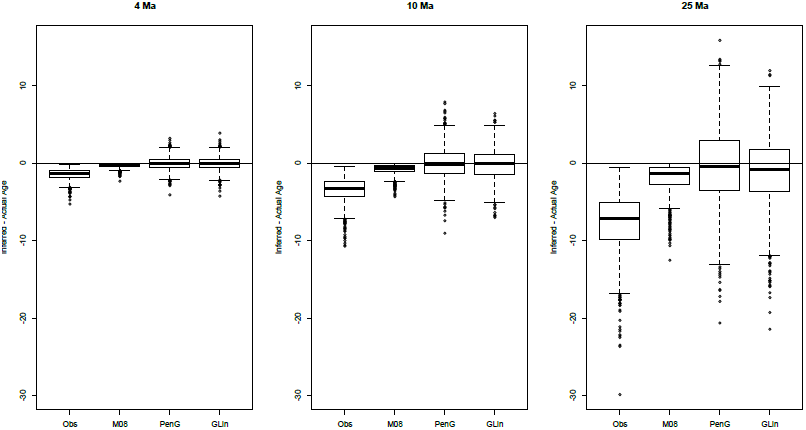
Mean bias in different methods when calibrating the true tree. Bias is calculated as actual age subtracted from inferred age. Results for three rates of fossilization are shown: every 4 Ma, 10 Ma, and 25 Ma. Solid line indicates median, box outlines 50% of samples, whiskers encompass 1.5 times the interquartile range above and below the upper and lower quartiles, and circles indicate outliers outside of this range. All methods underestimate divergence times with Obs being the most extreme case, particularly when fossil sampling is infrequent (25 Ma).

**Table 1.** Bias in age estimates when the true tree is calibrated using the methods shown. Bias is calculated as the actual divergence time subtracted from the estimated divergence time. Negative values indicate under-estimates; positive values indicate over-estimates. Results are ranked by the absolute value of the mean, with lower values indicating less bias. P-values testing for bias are based on the Wilcoxon signed rank test: bold p-values (p < 0.05) indicate that the hypothesis of an unbiased method can be rejected. The final row shows instances where the post-hoc Friedman’s test indicates that a method yields results significantly older than the method listed (p < 0.05; or † indicating p < 0.001). Because all means are negative, older results represent less biased estimates.

|  | Obs | M08 | PenG | GLin |
| --- | --- | --- | --- | --- |
| <u>Every 4 Ma</u> |  |  |  |  |
| Rank | 4 | 3 | 1 | 2 |
| Mean | -1.506 | -0.339 | -0.068 | -0.092 |
| St. Dev | 0.689 | 0.295 | 0.880 | 0.887 |
| Unbiased | **< 0.001** | **< 0.001** | **< 0.001** | **< 0.001** |
| Signif. older than |  | Obs† | Obs†<br>M08†<br>GLin† | Obs†<br>M08† |
| <u>Every 10 Ma</u> |  |  |  |  |
| Rank | 4 | 3 | 1 | 2 |
| Mean | -3.518 | -0.781 | -0.152 | -0.196 |
| St. Dev | 1.671 | 0.738 | 2.110 | 2.073 |
| Unbiased | **< 0.001** | **< 0.001** | **< 0.001** | **< 0.001** |
| Signif. older than |  | Obs† | Obs†<br>M08†<br>GLin | Obs†<br>M08† |
| <u>Every 25 Ma</u> |  |  |  |  |
| Rank | 4 | 3 | 1 | 2 |
| Mean | -7.886 | -1.886 | -0.716 | -1.240 |
| St. Dev | 3.855 | 2.002 | 4.828 | 4.640 |
| Unbiased | **< 0.001** | **< 0.001** | **< 0.001** | **< 0.001** |
| Signif. older than |  | Obs† | Obs†<br>M08†<br>GLin† | Obs†<br>M08† |

The pattern of bias shifts slightly for post-hoc calibration of an inferred tree (inferred from sequence data), as indicated in Figure 3 and Table 2. The PenG and GLin methods again show the least bias, though the slight underestimation bias is significant (Table 2). The Obs method continues to exhibit substantial underestimation bias across fossilization rates, and to produce significantly different results from all other methods (Table 2). However, at both the medium and high fossilization rates, the M08 method exhibits a significant **overestimation** bias, with mean bias significantly different from all other methods. In the Discussion, we explain why the M08 method is expected to give an overestimation bias when there is uncertainty in the inferred tree.

**Figure 3.**
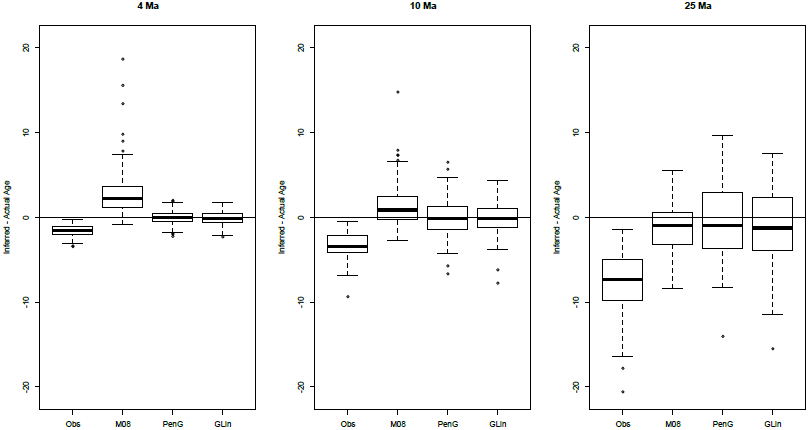
Mean bias in different methods for post-hoc calibration of a tree inferred from simulated sequence data. Results for three rates of fossilization are shown: every 4 Ma, 1 0 Ma, and 25 Ma. Error bars are as described in the legend to Fig. 2. The introduction of error from molecular data leads M08 to exhibit a bias toward overestimating divergence dates when fossil sampling is dense (i.e. 4 Ma and 10 Ma).

**Table 2.** Bias in age estimates when an ultrametric tree is inferred using simulated sequence data, then (separately) calibrated *post hoc* using the methods shown. See Table 1 for explanation of statistical tests and their interpretation. Because not all means have the same sign, older results do not necessarily indicate less biased estimates.

|  | Obs | M08 | PenG | GLin |
| --- | --- | --- | --- | --- |
| <u>Every 4 Ma</u> |  |  |  |  |
| Rank | 3 | 4 | 1 | 2 |
| Mean | -1.616 | 2.639 | -0.255 | -0.293 |
| St. Dev | 0.846 | 3.180 | 1.011 | 0.984 |
| Unbiased | **< 0.001** | **< 0.001** | **< 0.001** | **< 0.001** |
| Signif. older than |  | Obs†<br>PenG†<br>GLin† | Obs† | Obs† |
| <u>Every 10 Ma</u> |  |  |  |  |
| Rank | 4 | 3 | 1 | 2 |
| Mean | -3.531 | 1.345 | -0.317 | -0.456 |
| St. Dev | 1.674 | 2.511 | 2.166 | 2.081 |
| Unbiased | **< 0.001** | **< 0.001** | **0.004** | **< 0.001** |
| Signif. older than |  | Obs†<br>PenG†<br>GLin† | Obs† | Obs† |
| <u>Every 25 Ma</u> |  |  |  |  |
| Rank | 4 | 3 | 1 | 2 |
| Mean | -7.759 | -1.158 | -0.847 | -1.083 |
| St. Dev | 3.364 | 2.987 | 4.546 | 4.480 |
| Unbiased | **< 0.001** | **< 0.001** | **< 0.001** | **< 0.001** |
| Signif. older than |  | Obs† | Obs† | Obs† |

Figure 4 and Table 3 show the results of the test comparing the new Bayesian methods for inferring a calibrated tree in BEAST (PenG and GLin), with suitably adapted versions of Obs, Marshall (2008), and Dornburg, et al (2011). As with earlier tests, the Obs method exhibits significant and substantial underestimation bias. This approach is considerably less accurate than all other methods (Table 4) and this difference is significant in all comparisons, except against the aD11 method at low and medium fossil sampling. The M08min and aD11 methods exhibit strong overestimation bias at dense fossil sampling but perform well at sparse fossil sampling.

**Figure 4.**
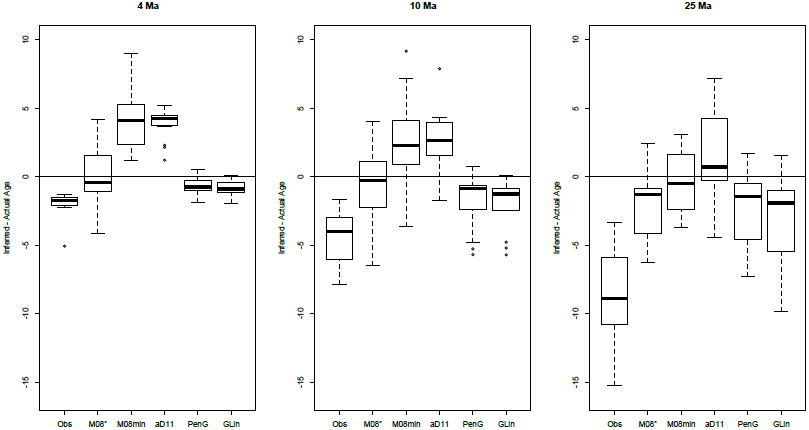
Mean bias in different methods of calibration implemented as prior probability distributions in BEAST. Results for three rates of fossilization are shown: every 4 Ma, 1 0 Ma, and 25 Ma. Error bars are as described in the legend to Fig. 2. The M08min and aD11 methods are biased toward overestimating divergence times when fossil sampling is dense (i.e. 4 Ma and 10 Ma). Obs, PenG, and GLin exhibit underestimation bias at all sampling densities, while M08 underestimates when fossil sampling is infrequent (25 Ma).

**Table 3.**
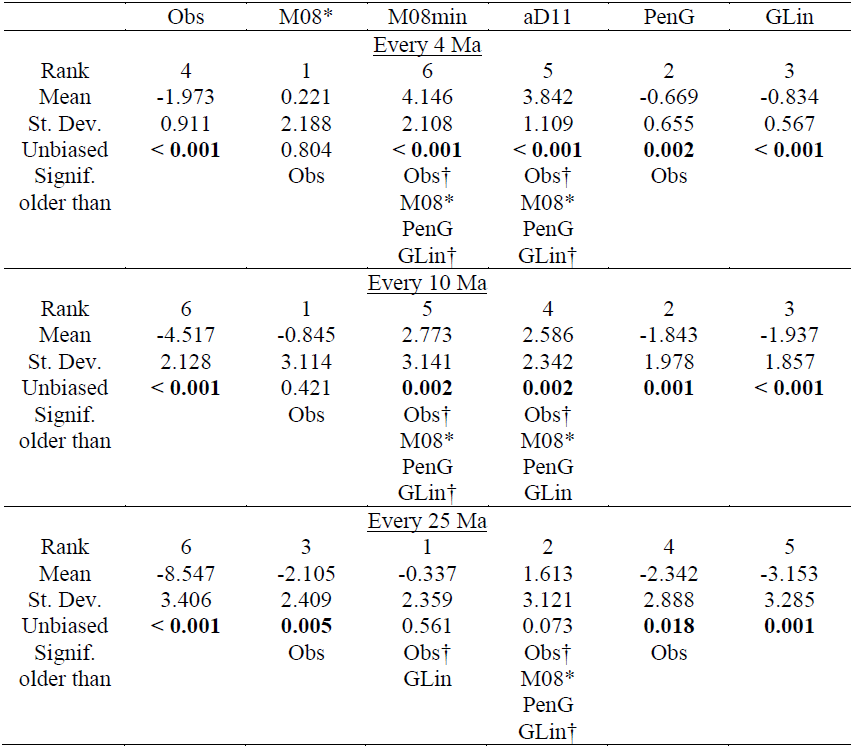
Bias in age estimates when a calibrated tree is inferred from sequence data while simultaneously incorporating fossil-based constraints or priors. See Table 1 for explanation of statistical tests and their interpretation.

The M08^*^ approach shows less bias than PenG and GLin, but this difference is not significant, and comes at the expense of a much wider spread of inferred values. Thus it is valuable to compare, not merely bias, but accuracy (closeness of an estimate to the truth) and precision (the width of the estimate), shown in Table 4 and Figure 5. When accuracy is measured as the count of nodes for which the 95 % highest posterior density (HPD) includes the true age, PenG and GLin perform the best, although not significantly better than M08^*^ and M08min. When precision is defined as the mean length of 95 % HPDs that include the true value, PenG and GLin are the most precise methods at all fossil densities (Figure 5), and are significantly more precise than M08^*^ (Table 4). We did not test the precision of the Obs method because there were too few instances of the actual value falling within the 95 % HPD.

**Table 4.** Accuracy and precision in age estimates when Bayesian analysis is calibrated with explicit priors. Accuracy is measured as a count of nodes (out of 7) where the 95 % HPD estimate includes the actual age. Instances where method yields a significantly higher count of accurate nodes based on the post-hoc Friedman’s test (with p < 0.05 and † indicating p < 0.001) than method named are shown. The % accurate nodes row indicates the total percentage of nodes which fall within the 95 % HPD. Precision is measured as the length of 95 % HPD ranges for those instances where the actual age is included in 95 % HPD estimate. Instances where method yields a significantly shorter 95 % HPD length based on Tukey’s HSD test (with p < 0.05 and † indicating p < 0.001) than method named are shown.

|  | Obs | M08* | M08min | aD11 | PenG | Glin |
| --- | --- | --- | --- | --- | --- | --- |
| <u>Every 4 Ma</u> |  |  |  |  |  |  |
| Rank (Accuracy) | 6 | 3 | 4 | 5 | 1 | 2 |
| Mean nodes recovered | 0.200 | 5.333 | 5.133 | 2.733 | 5.733 | 5.667 |
| St. Dev. | 0.414 | 0.724 | 0.834 | 1.387 | 1.100 | 1.234 |
| Signif. more than |  | Obs† | Obs† |  | Obs† | Obs† |
|  |  | aD11 | aD11 |  | aD11† | aD11 |
| % Accurate nodes | 2.9 % | 76.2 % | 73.3 % | 39.0 % | 81.9 % | 81.0 % |
| Rank (Precision) | - | 5 | 4 | 3 | 1 | 2 |
| Mean HPD length | - | 22.978 | 15.236 | 6.023 | 4.647 | 4.924 |
| St. Dev. | - | 10.256 | 5.621 | 2.526 | 1.079 | 1.554 |
| Signif. shorter than | - |  | M08*† | M08*† | M08*† | M08*† |
|  |  |  |  | M08min† | M08min† | M08min† |
| <u>Every 10 Ma</u> |  |  |  |  |  |  |
| Rank (Accuracy) | 6 | 3 | 4 | 5 | 2 | 1 |
| Mean nodes recovered | 0.800 | 5.333 | 5.133 | 4.400 | 5.467 | 5.600 |
| St. Dev. | 1.0823 | 0.617 | 0.640 | 0.737 | 1.506 | 1.549 |
| Signif. more than |  | Obs† | Obs† |  | Obs† | Obs† |
|  |  |  |  |  | aD11 | aD11 |
| % Accurate nodes | 11.4 % | 76.2 % | 73.3 % | 62.9 % | 78.1 % | 80.0 % |
| Rank (Precision) | - | 5 | 4 | 3 | 1 | 2 |
| Mean HPD length | - | 24.450 | 17.792 | 11.528 | 7.886 | 8.031 |
| St. Dev. | - | 11.999 | 8.420 | 8.689 | 2.232 | 2.449 |
| Signif. shorter than | - |  |  | M08*† | M08*† | M08*† |
|  |  |  |  |  | M08min | M08min |
| <u>Every 25 Ma</u> |  |  |  |  |  |  |
| Rank (Accuracy) | 6 | 4 | 4 | 2 | 3 | 1 |
| Mean nodes recovered | 1.200 | 5.133 | 5.133 | 5.467 | 5.267 | 5.667 |
| St. Dev. | 0.941 | 0.743 | 0.743 | 1.125 | 1.624 | 1.291 |
| Signif. more than |  | Obs | Obs | Obs† | Obs† | Obs† |
| % Accurate nodes | 17.1 % | 73.3 % | 73.3 % | 78.1 % | 75.2 % | 81.0 % |
| Rank (Precision) | - | 5 | 4 | 3 | 1 | 2 |
| Mean HPD length | - | 22.497 | 15.753 | 13.817 | 13.525 | 13.778 |
| St. Dev. | - | 12.145 | 5.158 | 5.536 | 4.385 | 4.512 |
| Signif. shorter than | - |  | M08* | M08*† | M08*† | M08*† |

**Figure 5.**
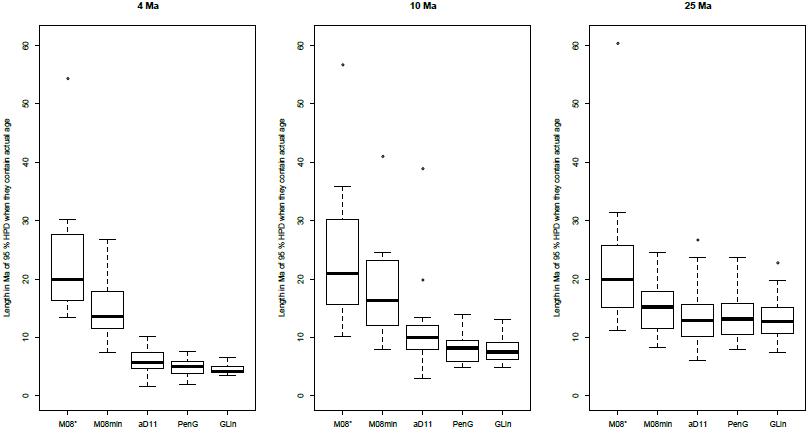
Accuracy and precision of different methods of calibration implemented as prior probability distributions in BEAST. Results for three rates of fossilization are shown: every 4 Ma, 1 0 Ma, and 25 Ma. Error bars are as described in the legend to Fig. 2. Precision is calculated as the mean length of 95 % HPD ranges that include the actual age. Methods which employ multiple calibrations (aD11, PenG, and GLin) generate smaller 95 % HPD intervals than those relying on a single calibration (M08^*^) or a single calibration plus minimum constraints (M08min).

The approach of using post-hoc calibrations was intended to identify biases outside of a full Bayesian analysis that, as shown by Heled and Drummond (2012), can be subject to complex effects due to calibration priors interacting with each another and with the Yule prior. We executed several sample BEAST runs with empty alignments and confirmed that many replicates within our simulations that employed PenG, GLin, and aD11 methods showed evidence that calibrating priors were interacting with one another even in the absence of genetic data. In spite of this interaction, our PenG and GLin simulations yielded reasonably accurate results, with 75 % to 82 % of the node ages correctly recovered within the 95 % HPD values.

## Discussion

Several generalities emerge from the comparative evaluation of methods above. Use of the oldest fossil for each node without correction (Obs method) suffers from a substantial underestimation bias across all fossil sampling rates, for all 3 tests— unsurprising given that fossils are inherently underestimates of true divergence times.

The M08 approach of Marshall (2008) represents a genuine improvement. When the true tree is calibrated with fossils post-hoc, M08 performs with little bias (Fig. 2). Ages are underestimated, but the magnitude of this underestimate is significantly smaller than the Obs method. This underestimation is consistent with Marshall’s (2008) expectation, and he provided a method to calculate an upper 95 % estimation. We did not evaluate the performance relative to that upper confidence interval, but have every expectation that it would encompass the true value at all or most nodes for the true tree test.

As noted above, the reason for carrying out separate post-hoc calibrations with a true tree and an inferred tree is to tease out different sources of bias. In the case of Marshall’s method, the comparison of these two tests reveals an interaction with tree inference that transforms an under-estimation bias (Fig 2, Table 1) into an overestimation bias (Fig 3, Table 2), a result consistent with Lukoschek et al. (2012). This effect may be explained as follows, as a consequence of tree inference error. Consider a case involving an ideal set of fossil calibrations— for each clade, a fossil is present that is arbitrarily close (e.g., 1 year) to the true time of origin—, and an inferred molecular phylogeny with the correct topology, but with branch lengths subject to error. In the inferred ultrametric tree, the relative ages of some nodes will be too high, and others too low; there will be a node with the greatest upward error relative to the root, and another with the greatest downward error. Although the fossil record is ideal, such that no fossil is better than any other, M08 entails picking the single fossil calibration that yields the oldest tree. In such a case, M08 will calibrate the tree using the node with the greatest downward error in placement: the age of the calibrated node will be correct (given the ideal fossil record), but the ages of all other nodes will be over-estimates. More generally, in real cases where the fossil record is not perfect, M08 will tend to err by calibrating on a node with a downward placement error.

The over-estimation bias disappears in the third set of tests for M08^*^, but the comparison with M08min indicates that the over-estimation effect is masked by another effect. The 95 % HPD values produced in the M08^*^ analyses include ages at nodes that are younger than the minima indicated by excluded fossil calibrations at those nodes. This situation reflects our choice to adapt M08 for a Bayesian approach by calibrating a single MCC tree (namely, the best tree), instead of by calibrating each tree in the posterior sample from the Markov chain. Marshall (2008) explicitly designed his approach to avoid the potential that age estimates are younger than the fossil record allows. This situation is prevented in M08min by adding all fossils as hard minima, and the use of this modification causes the over-estimation bias to re-emerge (Fig 4).

The aD11 approach also suffers from an overestimation bias. The 95 % HPD estimates included significantly fewer actual node ages than all approaches except the Obs method. By including a lognormal prior shape, the approach of Dornburg et al. (2011) appears to be magnifying the overestimation bias already inherent to M08. We note again that aD11 is an abbreviated (less computationally burdensome) implementation of the approach of Dornburg and colleagues (2011): limited tests verify that the abbreviated approach behaves the same (see Methods), but we do not claim to have tested precisely the method of Dornburg, et al.

Possibly a large fraction of published analyses suffer from either the underestimation bias inherent to the Obs method, or the overestimation bias that affects M08 and related methods under the condition of error in tree inference. Although explicit use of Marshall’s (2008) method has been limited (Davis et al. 2009; Aschliman et al. 2012; Miya et al. 2013; Yang et al. 2013; Dornburg et al. 2014; Hoekzema and Sidlauskas 2014; Kenaley et al. 2014; Kranitz et al. 2014), the underlying philosophy may be far more widespread. Brochu et al. (2004) suggest that many potential fossil calibrations are thrown out in the early stages of studies on the grounds of yielding anomalously young ages at other nodes with fossils. Our study suggests that this practice has the potential to bias age estimates toward older dates, particularly when the fossil record is good. By using a method focused on correcting for the well-established underestimation bias that results from using observed fossils as point estimates, researchers may be converting errors in branch length inference based on molecular data into an overestimation bias.

Compared to other approaches to specifying constraints or Bayesian priors while inferring a scaled tree, PenG and GLin achieve better accuracy with higher precision, exhibiting smaller HPD intervals around age estimates (Fig. 5, Table 4). Both methods exhibit a small but significant underestimation bias that is worse for sparse fossil sampling. This bias may arise from the particular way we implement PenG and GLin. When fossils are sparse in a clade, such that there is not a pair of interval-defining fossils on its deepest branches, we simply descend into subclades to define an interval, in order to increase precision by maximizing the number of calibrations available. When intervals are defined in this way, by descending into a sub-clade, the relevant rate of fossil recovery in our simulations is not constant, but accelerating (because the rate along a lineage doubles when it splits), with the result— a violation of our stated model— that the expected length of the interval from the origin of the clade to the first fossil (UltG) is greater than the expected length of PenG or GLin/2. Consequently, to the extent that sparse fossil sampling results in intervals not on the deepest branches, PenG and GLin/2 will be under-estimates of UltG, and the age of the node will be underestimated. Presumably this bias would disappear, at the cost of some precision, if PenG and GLin were implemented more strictly to rely only on the deepest branches.

The approaches to calibration and evaluation used here involve other simplifying assumptions whose impact on estimation using naturally evolved data is difficult to predict. For instance, our evaluation scheme includes no error in the assignment of fossils to clades, whereas in reality this is an important issue. Likewise, our evaluations use a simple number for fossil ages, whereas actual fossil ages carry uncertainty. Our simulations assume that fossil recovery is independent across branches, which is unlikely to be true when dealing with real data (Wagner and Marcot 2013). Likewise, gaps and other non-uniformities exist in the fossil record across time and space. However, Solow (2003) argues that PenG remains a good estimator for UltG in the case of heterogeneity, on the grounds that the conditions associated with deposition of the oldest and second-oldest fossils are likely to be similar. That is, the assumption of homogeneity in interval-based methods is a local, not a global, assumption.

Our methods are not the only available methods based on a prior that can be justified. Nowak et al. (2013) have developed an approach for generating well-justified prior shapes for calibrating nodes, but their method, which is based on Foote (2000), requires a considerable amount of information such as known fossil diversity, first and last occurrence dates of fossil taxa, estimates of extant diversity, and estimates of preservation and recovery of fossils. As is the case for tip-dating approaches, the information necessary to apply the method of Nowak et al. (2013) is unlikely to be available for most phylogenetic studies. Using a method that requires even more information, Wilkinson et al. (2011) also introduced an approach for calibrating prior shapes and applied it to a well-studied primate dataset. Avoiding the need for calibration prior shapes, the fossilized birth-death method of Heath et al. (2014) works like a tip-dating method but does not require a morphological dataset. Their method models macroevolutionary parameters in the absence of an explicit calibration prior and can be used in instances where only limited fossil information is present. Nevertheless, Heath et al (2014), Nowak et al. (2013), and Wilkinson et al. (2013) require the modeling of a number of phylogeny-wide parameters such as speciation and extinction rates even though these parameters are likely to vary not only over time, but may specifically differ early versus late in the evolutionary history of a successful lineage. By focusing on only the first few fossils along a lineage, our approaches minimize the assumption that aspects of preservation, recovery, speciation, and extinction have remained unchanged over time (Solow 2003).

## Conclusions

In the work described here, we have introduced two approaches to Bayesian calibration based on fossil intervals. We have specified conditions under which PenG and GLin/2 are point estimates of an important unknown quantity— UltG, the time from the oldest fossil to the root of a clade. Furthermore, assuming an uninformative and arguably optimal (Jeffreys) prior for the unknown rate of fossilization, we show that, given the value of either PenG or Glin/2, the corresponding posterior distribution for UltG is log-logistic. When tested within a Bayesian framework for inferring a scaled tree from simulated sequence and fossil data, the PenG and GLin methods are considerably more precise than the other methods tested, without significantly greater bias, entailing an overall increase in the accuracy of inferred divergence times (Figs. 4 and 5, Tables 3 and 4). When tested as methods to provide point estimates of node ages in post-hoc calibrations using simulated data, PenG and GLin exhibit less bias than other methods tested (Figs. 2 and 3, Tables 1 and 2).

The work described here establishes a rigorous basis, previously missing, to specify an informed prior on the age of a node, useful for Bayesian analyses that employ multiple calibration points. The new methods can be implemented in BEAST by specifying appropriate parameters in the input file (see Methods for details). These new methods do not require extensive fossil data: they may be applied to actual datasets when one knows the oldest fossil along with either the next-oldest fossil, or the oldest fossil on a sister lineage. Both methods are based on a very general property of Poisson processes (with other possible applications), namely that successive events from a process define an interval that is informative about the lengths of adjacent intervals.

### Availability of supporting data

The data sets supporting the results of this article are available in the GitHub repository, http://github.com/arlin/PenG_and_GLin.

## List of abbreviations

aD11: – An abbreviated version of the method of Dornburg et al. (2011).
GLin: –Ghost Lineage Length, the difference between the oldest fossil on one lineage, and the oldest fossil on a sister lineage. The GLin method uses GLin/2 in approximating the actual age of a node.
M08: – Marshall’s (2008) method.
M08^*^: – Adaptation of Marshall’s (2008) method for a Bayesian context.
M08min: – Adaptation of Marshall’s (2008) method for Bayesian context, preventing age estimates for a node that are more recent than known fossils at that node.
Obs: – Observed age, the oldest known fossil belonging to a clade. The Obs method uses this value without correction as an estimate of the actual age of a node.
PenG: – Penultimate Gap, the difference between the oldest fossil in a clade and the second oldest fossil. The PenG method uses this value when approximating the actual age of a node.
UltG: – Ultimate Gap. The difference between the oldest known fossil and the actual origin of a clade.

## Competing interests

The authors declare that they have no competing interests.

## Authors’ contributions

RWN designed and carried out the study, performed statistical analyses, and drafted the manuscript.

CLS Modified indel-Seq-Gen to incorporate fossilization events along the phylogeny, generated simulated data sets.

DMM contributed analytical tools and helped to draft the manuscript.

AS participated in the design of the study and helped to draft the manuscript.

All authors read and approved the final manuscript.

## Acknowledgments

A. L. Rukhin provided valuable mathematical assistance. C. W. Kilpatrick and B. Akaogi were part of early discussions leading to the approaches presented here. P. J. Wagner, J. L. Thorne, R. Wilkinson, and T. J. Near provided valuable suggestions on an earlier version of the manuscript. This research was performed while RWN held a National Research Council Research Associateship Award at the National Institute of Standards and Technology. This work was also supported by the National Institutes of Health (GM090201 to CLS). Certain commercial equipment, instruments, or software are identified in this paper to foster understanding. Such identification does not imply recommendation or endorsement by the National Institute of Standards and Technology, nor does it imply that the products identified are necessarily the best available for the purpose.

## Supplementary Figure

**Figure S1.** Accuracy of the lognormal approximation of the log-logistic distribution. BEAST does not implement a log-logistic prior, therefore we calculated the shape parameters to approximate it using the lognormal. The two distributions are compared here. The gray line is the probability density function for a random variable UltG/*x*, where UltG is distributed according to its posterior distribution given that the observed value for PenG is *x*. The black line is our log-normal approximation, which was constructed as described in the main text. The error in the log-normal approximation is no more than 24 % over the range 1/100 < UltG/x < 100. The figure would be identical if we instead defined *x* to be the observed value of GLin/2.

